# Dysfunction of the histone demethylase IBM1 in Arabidopsis reshapes the root-associated bacteria community

**DOI:** 10.1101/2020.10.19.344762

**Authors:** Suhui Lv, Yu Yang, Li Peng, Kai Tang, Richa Kaushal, Huiming Zhang

## Abstract

Root-associated bacteria communities are influential to plant growth and stress tolerance. Epigenetic regulation plays important roles in many plant biological processes, but its potential impacts on the assembly of root microbiota remain unclear. Here we report that dysfunction of the histone demethylase IBM1 in Arabidopsis substantially alters root-associated soil bacteria community. We compared two alleles of *ibm1* mutant (*ibm1-1* and *ibm1-4*) with wild type Arabidopsis regarding the root-associated bacteria community by 16S rRNA gene sequencing. The constrained principal coordinate analysis (PCoA) showed that the *ibm1* mutants are both clearly separated from Col-0 along the major coordinates. Among the 29 families which have a relative abundance more than 0.5% in at least one sample, 10 and 11 families were commonly affected by *ibm1-1* and *ibm1-4* alleles in the rhizosphere and the endosphere compartment, respectively. Notably, the ACMs (Abundant Community members) belonging Pseudomonadaceae showed increased relative abundance in the *ibm1* mutant alleles compared to Col-0 in both the rhizosphere and the endosphere compartments. The ACMs belonging to Oxalobacteraceae mostly showed decreased relative abundance in *ibm1* mutants compared to Col-0 in endosphere compartment. These findings demonstrate an influential function of IBM1-mediated epigenetic regulation in shaping the root-associated microbiota.

## Introduction

The plant rhizosphere, which refers to a thin layer of soil adhering to the roots, harbors various microorganisms especially bacteria in terms of species richness (1). While some rhizobacteria have no observable effects on plants, others are either pathogens that cause detrimental effects on plants or beneficial strains termed plant growth-promoting rhizobacteria (PGPR) that promote plant vigor (1, 2). The root-associated bacteria community has been shown to be important for plants health (3), and is therefore emerging as an important target for soil management and for studying plant-microbe interactions.

The plant immune response plays an important role in sculpting the plant-associated microbiome. Evidences have shown that plant-associated microbiome can be changed when plants are attacked by pathogen or insect (3–5). For example, foliar pathogen *Pseudomonas syringae* pv tomato (Pst DC3000) induces the Arabidopsis root secretions of L-malic acid (MA), which selectively recruits the beneficial rhizobacterium *Bacillus subtilis* FB17 (6). For another example, aphid (Myzus persicae) infestation on pepper plants increased the root population of the beneficial *Bacillus subtilis* GB03, but reduced population of the pathogenic *Ralstonia solanacearum* SL1931 (7). Thus, it seems that plants are able to recruit beneficial microorganisms in the rhizosphere to help suppress pathogens through immune response upon attacked. Besides, analysis of Arabidopsis root microbiota using mutants which either constitutively accumulates defense phytohormone salicylic acid (SA) or impaired in SA accumulation/signaling have revealed that SA modulates colonization of the root by specific bacterial families (8). Moreover, the Arabidopsis quadruple mutant (*min7 fs2 efr cerk1*; hereafter, *mfec*), simultaneously defective in pattern-triggered immunity and the MIN7 vesicle-trafficking pathway, also harbors altered endophytic phyllosphere microbiota (9). These findings demonstrate that plant immunity system functions in shaping plant associated microbiome.

In the past decade, the role of epigenetic factors in controlling plant immunity has been emerging. Several studies have implicated that DNA demethylation functions in plant immune response (10–14). For example, the triple DNA demethylase mutant, *rdd* (*ros1 dml2 dml3*), shows increased susceptibility to the fungal pathogen *Fusarium oxysporum*, accompanied with decreased expression of many stress response genes(11). Further investigations indicate that DNA demethylase target promoter transposable elements to positively regulate stress-responsive genes (11). Meanwhile, RdDM component *nrpe1* displayed enhanced resistance to the biotrophic pathogen *Hyaloperonospora arabidopsidis* (*Hpa*), which is associated with elevated SA-dependent PR1 gene expression upon *Hpa* infection(12). Moreover, global disruption of DNA methylation by *met1* or *ddc* mutation activates defense responses against *P. syringae* (13). Interestingly, plants exposed to bacterial pathogen, avirulent bacteria, or SA hormone showed numerous stress-induced differentially methylated regions, many of which were intimately associated with differentially expressed genes (13). Some TEs are demethylated and transcriptionally reactivated during antibacterial defense in Arabidopsis, and DNA demethylation restricts multiplication and vascular propagation of the bacterial pathogen *Pseudomonas syringae* in leaves (14). These evidences indicate that DNA demethylation acts as part of the plant immune response. Besides DNA demethylation, histone modifications and chromatin remodeling also take part in plant immunity control. For example, an Arabidopsis JHDM2 (JmjC domain-containing histone demethylase 2) family protein JMJ27, which displays H3K9me1/2 demethylase activity, is induced in response to virulent *Pseudomonas syringae* pathogens and is required for resistance against these pathogens (15). For another example, H2A.Z is the variant of the canonical histone H2A, and SWR1-like complex is responsible for the H2A.Z deposition (16).

Mutation of H2A.Z coding genes (*hta9 hta11*) or SWR1 complex components (pie1, sef) caused constitutive expression of systemic acquired resistance (SAR) marker genes and increased resistance to the virulent *Pseudomonous syringae* (16). Altogether, it is evident that epigenetic regulation, including DNA (de)methylation, histone modification, and chromatin remodeling, plays an important role in plant immune response.

Arabidopsis IBM1 (increase in bonsai methylation 1) encodes a Jmjc-domain containing H3K9 demethylase, mutation of which resulted in ectopic H3K9me and non-CG methylation in thousands of genes, as well as a variety of developmental phenotypes (17–20). The *ibm1*-induced genic H3K9me2 depends on both histone methylase KYP/SUVH4 and DNA methylase CMT3 (20). Later it was found that developmental defects of *ibm1* can be suppressed by a second mutation in *LDL2* (*LYSINE-SPECIFIC DEMETHYLASE 1-LIKE 2*) gene, a homolog of H3K4 demethylase (21). Interestingly, the *ldl2* mutation suppressed the developmental defects without suppressing the *ibm1*-induced ectopic H3K9me2 (21). It was proposed that the ectopic H3K9me2 mark directed removal of gene-body H3K4me1 and caused transcriptional repression in an LDL2-dependent manner (21). In another study, it was shown that the *ibm1* mutation caused hypermethylation of H3K9 and DNA non-CG sites within RDR2 and DCL3, which are two components in the RNA-directed DNA methylation (RdDM) pathway. The down-regulation of RDR2 and DCL3 gene expression in *ibm1* affected siRNA biogenesis in a locus-specific manner and disrupted RdDM-directed gene repression. (22). Thus dysfunction of IBM1 has a strong impact on plant epigenetic landscape including histone modifications and DNA methylation patterns.

In this study, we compared two alleles of *ibm1* mutant with wild type Arabidopsis regarding the root-associated bacteria community. The 16S rRNA gene sequencing results revealed substantial impacts of the *ibm1* mutations on the bacteria communities in the rhizosphere and endosphere. Particularly, we show that the Pseudomonadaceae family is enriched in the *ibm1* mutant alleles compared to Col-0 in both rhizosphere and endosphere compartments, while the Oxalobacteraceae family showed decreased relative abundance in *ibm1* mutants compared to Col-0 in endosphere compartment. Our study of IBM1 demonstrates an influential function of epigenetic regulation in shaping the root-associated microbiota.

## Results

### *Ibm1* mutation substantially alters plant root-associated microbiota

To evaluate the potential effects of *ibm1* mutation on the plant-bacteria interaction, we investigated the rhizosphere and root endosphere microbiomes of Col-0 and two *ibm1* mutant lines (*ibm1-1* and *ibm1-4*). We collected field soil from Chenshan Botanical Garden, Shanghai, China, and grew the Col-0, *ibm1-1* and *ibm1-4* in this field soil. 17 days later, the DNA were extracted from bulk soil, rhizosphere and endosphere compartments of different genotypes. Then the bacterial 16S rRNA gene amplicon libraries were generated with the PCR primers 799F and 1193R harboring the hypervariable regions V5-V6-V7 (23, 24), and the bacteria communities were profiled by 16s rDNA sequencing.

A total of 1,847,563 high-quality sequences were obtained with a median read count per sample of 57,784 (range 7,511-137,080) across all the 35 samples. To correct the sequencing depth differences among samples, these high-quality sequences were further subjected to rarefaction to 7000 sequences per sample based on the rarefaction curve (Figure 1A), which resulted in 4688 unique OTUs as the threshold-independent community (TIC) (Dataset S1). A minimum of 20 sequences per OTU in at least one sample was used as a criterion to define abundant community members (ACMs). The ACM of all samples was represented by 186 bacterial OTUs comprising of 66.73% of the rarefaction quality sequences (Dataset S1). We normalized this table by dividing the reads per OTU in a sample by the sum of the usable reads in that sample, resulting in a table of relative abundances (RA) (Dataset S1) for quantitative comparison.

**Figure 1.**
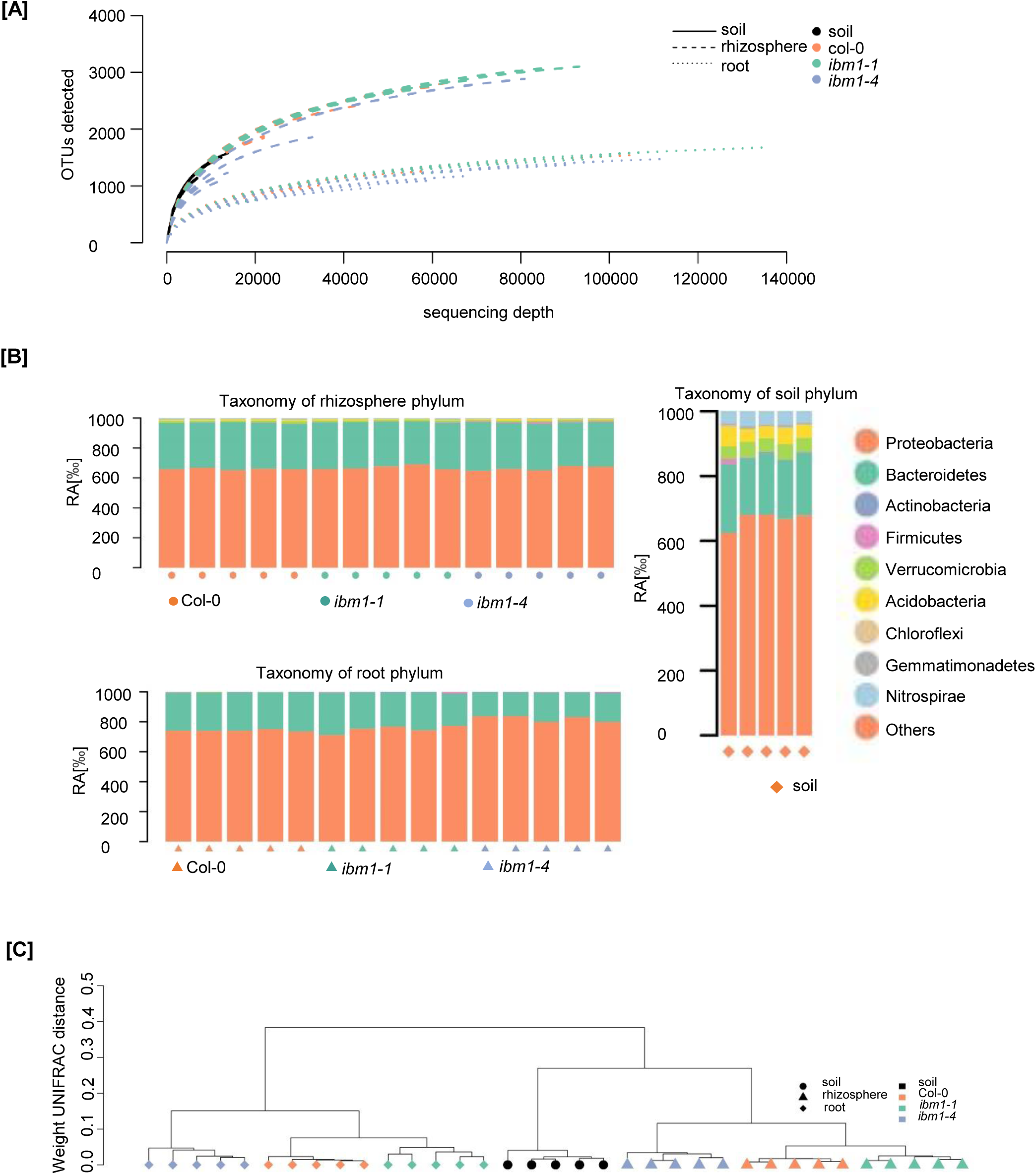
**[A]** Rarefaction analysis. The pooling of the sequences from all the samples was permitted to display the bacterial communities with a sequencing depth of 7,511 – 137,080 quality sequences per sample. **[B]** Taxonomic structure at the phylum rank of the ACM. **[C]** Beta diversity. Between-sample diversity was calculated for ACM by weighted UniFrac distance metric on 7,000 sequences per sample.

We first examined taxonomic composition in the whole dataset. The bulk soil samples are mainly composed of Proteobacteria and Bacteroidetes, with smaller portions of Acidobacteria (4.6%), Verrucomicrobia (4.4%), Nitrospirae (4.0%), Gemmatimonadetes (0.61%), Firmicutes (0.40%), Actinobacteria (0.22%), and Chloroflexi (0.11%) (Figure 1B;Dataset S1). Meanwhile, the rhizosphere and root communities were both dominated by Proteobacteria and Bacteroidetes regardless of genotypes (Figure 1B). Then we used the weighted UniFrac metric to compare community diversity between samples (25) (Figure 1C). Consistent with previous studies (26-28), the hierarchical clustering of UniFrac distances revealed that the compartments (bulk soil, rhizosphere, endosphere) are the major sources of variation in bacteria communities.

Next, we performed constrained principal coordinate analysis (PCoA) of ACM to evaluate the impacts of *ibm1* mutation on bacteria communities in the rhizosphere and the endosphere. The *ibm1* mutants are both clearly separated from Col-0 along the major coordinates, which represent 79% and 73% variances for the rhizosphere and the endosphere, respectively (Figure 2A and B), indicating that IBM1 dysfunction alters the microbial communities in these two compartments.

**Figure 2.**
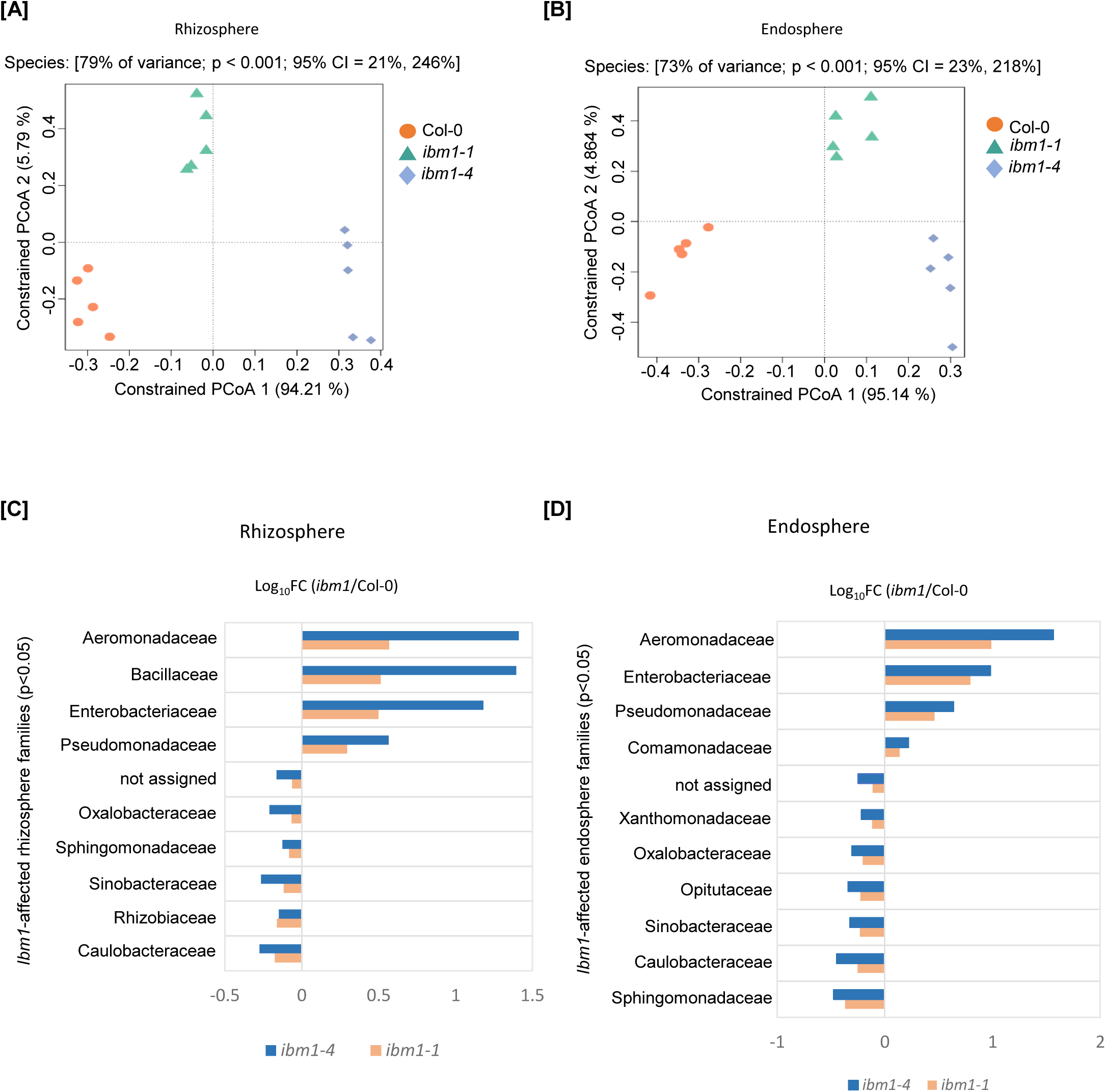
**[A]** and **[B]** Constrained principal coordinate analysis (PCoA) of ACM in the rhizosphere [A] and endosphere [B] of Col-0, *ibm1-1* and *ibm1-4*. The significant ACMs (fold change>2, FDR<0.05) were used for canonical analysis of principal coordinates, which was constrained for the genotypes. The percentage of variation explained by each axis refers to the fraction of the total variance of the data (ACM) explained by the constrained factor. **[C]** and **[D]** The families which are commonly affected by *ibm1-1* and *ibm1-4* (p<0.05) in rhizosphere [C] and endoshphere [D], separately. FC: fold change.

To some extent, the *ibm1-1* and *ibm1-4* alleles are also separated to each other along the major coordinates (Figure 2A and B), which may result from their different mutation types. Thus, we only consider those bacteria communities which were commonly affected in *ibm1-1* and *ibm1-4* as the “*ibm1*-affected” bacteria communities (P<0.05). At the phylum level, Firmicutes were enriched in both *ibm1-1* and *ibm1-4* rhizosphere compartments, and Verrucomicrobia were depleted in both *ibm1-1* and *ibm1-4* endosphere compartments (Dataset S1). However, only 1 ACM belonging to Firmicutes was identified as significantly enriched in *ibm1* mutants; and only 1 ACM belonging to Verrucomicrobia was identified as significantly depleted in *ibm1* mutants (Datset S1). Then we looked into the “*ibm1*-affected” bacteria communities at family level (Dataset S1, Figure 2C and D). In rhizosphere compartment, 10 of 29 families (RA > 5‰) were commonly affected in both *ibm1* mutant alleles (P<0.05), among which 4 are enriched and 6 are decreased in *ibm1* mutants (Figure 1C). The 4 families enriched in *ibm1* mutants include Pseudomonadaceae, Enterobacteriaceae, Bacillaceae, and Aeromonadaceae; while the 6 families with decreased relative abundance in *ibm1* mutants are Caulobacteraceae, Rhizobiaceae, Sinobacteraceae, Sphingomonadaceae, Oxalobacteraceae and a “not_assigned” family (Figure 2C; Dataset S1). In endosphere compartment, 11 families are commonly affected in both *ibm1-1* and *ibm1-4* alleles, among which 4 are enriched (Comamonadaceae, Pseudomonadaceae, Enterobacteriaceae, and Aeromonadaceae) in the *ibm1* mutants, and 7 showed decreased relative abundance (Sinobacteraceae, Oxalobacteraceae, Caulobacteraceae, Sphingomonadaceae, Xanthomonadaceae, Opitutaceae and a “not assigned” family) in *ibm1* mutants (Figure 2D; Dataset S1).

At the ACM level, among 164 ACMs with a relative abundance more than 5‰, 35 were commonly affected by *ibm1-1* and *ibm1-4* in rhizosphere compartment, while 36 were commonly affected by *ibm1-1* and *ibm1-4* in endosphere compartment (Dataset S1). In rhizosphere, 14 of the *ibm1*-affected ACMs are upregulated in the *ibm1* mutants, of which 6 belong to Pseudomonadaceae, 1 belongs to Aeromonadaceae, 3 belong to Comamonadaceae, 1 belongs to Flavobacteriaceae, 2 belong to Enterobacteriaceae and 1 belong to Bacillaceae (Figure S2A). Meanwhile, 21 of the *ibm1*-affected rhizosphere ACMs are downregulated in *ibm1* mutants, including ACMs belonging to Rizobeaceae (3), Flavobacteriaceae (3), Comamonadaceae (3), Xanthomonadaceae (1), Oxalobacteraceae (2), Sphingomonadaceae (1) Sinobacteraceae (1), Caulobacteraceae (2) and 5 ACMs with “not assigned” family (Figure S2B). In endosphere, 13 of the *ibm1*-affected ACMs are upregulated in the *ibm1* mutants, including 4 belonging to Pseudomonadaceae, 1 belonging to Aeromonadaceae, 5 belonging to Comamonadaceae, 1 belonging to Rizobeaceae, 1 belonging to Methylophilaceae, 1 belonging to Enterobacteriaceae (Figure S2C). 23 of the *ibm1*-affected ACMs are downregulated in the *ibm1* mutants, including 6 from Oxalobacteraceae, 6 from Comamonadaceae, 2 from Caulobacteraceae, 1 from Cytophagaceae, 1 from Opitutaceae, 1 from Rhizobiaceae, 1 from Sinobacteraceae, 1 from Sphingomonadaceae, and 4 with “not assigned” family (Figure S2D). Altogether, these results revealed an influential role of IBM1 in the assembly of plant-associated bacteria community.

Notably, in both rhizosphere and endosphere compartments, the ACMs (RA>5‰) belonging to the families of Pseudomonadaceae all showed increased relative abundance in the *ibm1* mutant alleles compared to Col-0 (Figure 3A and B; Dataset S1). Besides, in endosphere compartment, ACMs belonging to Oxalobacteraceae mostly showed decreased relative abundance in *ibm1* mutants compared to Col-0 (Figure 3C;Dataset S1). It was documented that Pseudomonadaceae family are enriched in the root microbiota of *cpr5* and *cpr1* mutants, which constitutively produce and accumulate SA contents; while Oxalobacteraceae family are enriched in the *pad4* mutant, which is impaired in SA signaling (8). Thus there is a possible correlation between high SA level and plant association with Pseudomonadaceae and Oxalobacteraceae.

**Figure 3.**
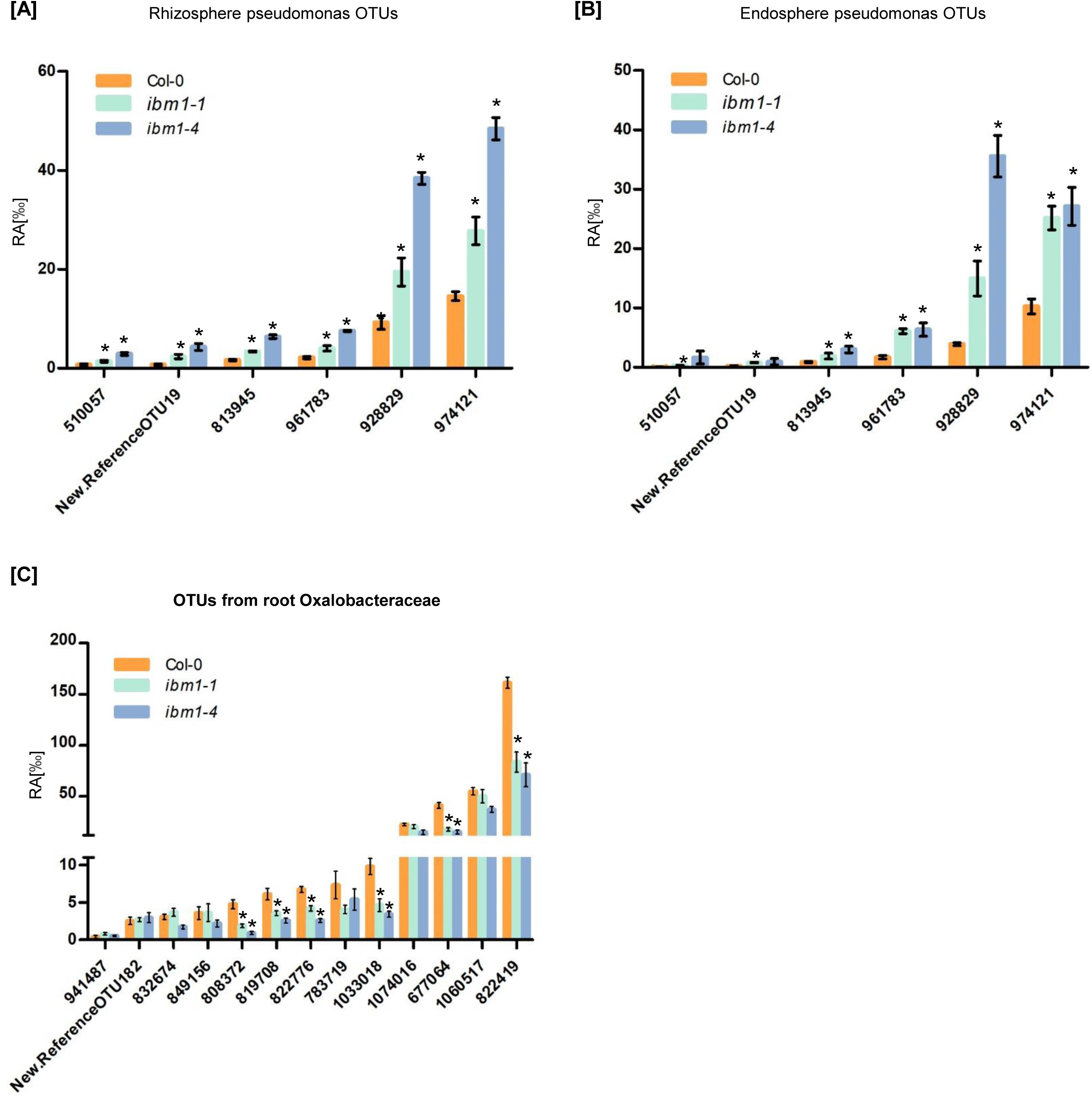
**[A]** and **[B]** Relative abundance (RA) of ACMs belonging to Pseudomonadaceae at rhizosphere [A] and endosphere [B], separately. **[C]** Relative abundance of ACMs belonging to Oxalobacteraceae at endosphere compartment. Values are mean ± SE, n=5. Asterisks indicate significant differences between *ibm1* mutant and wild type Col-0 (p < 0.05, ANOVA).

## Discussion

A distinct pattern in ibm1 microbiota is that the Pseudomonadaceae family are enriched in ibm1 rhizosphere and endosphere compartments, while Oxalobacteraceae family are depleted in the endosphere compartment. It was reported that SA modulates colonization of the root by specific bacterial families (8). From the published microbiota data, we found that Pseudomonadaceae family are enriched in the root microbiota of *cpr5* and *cpr1* mutants, which constitutively produce and accumulate SA contents; and Oxalobacteraceae family are enriched in the *pad4* mutant, which is impaired in SA signaling (8). These indicate a role of increased SA in affecting the Pseudomonadaceae and Oxalobacteraceae root colonization in *ibm1* mutant. In addition to SA, other factors probably also contribute to shaping the ibm1 root microbiota. For example, flavonoids and coumarins are known Arabidopsis root exudates (29) which may affect the root-associated microbiota. In future, the root exudates profiles of *ibm1* may be necessary to explain how the root-associated microbiota changed in *ibm1* mutant.

## Methods

### Plant material and growth condition

Soil from the field in Shanghai Chen Shan Botanical Garden was collected, homogenized and mixed with commercial soil (Pindstrup Substrate) with 2:1 ratio, and placed into 10×10×8cm pots. Then 14-day-old of Col-0, *ibm1-1* and *ibm1-4* seedlings were transplanted into the pots (five seedlings for each pot), and grown for another 17 days in a growth room at 22°C under 16h light/8h dark condition. 5 biological replicates were prepared for each genotype and each biological replicate contains 10 plants. Unplanted pots were subjected to the same conditions as the planted pots to prepare the control soil samples at harvest.

### DNA sample preparation

The rhizosphere samples and root endophyte samples were harvested according to Schlaeppi et al. (28). DNA were extracted from those samples by FastDNA® SPIN kit for Soil (MP Biomedicals). Samples were homogenized in the Lysis Matrix E tubes using a Retsch MM400 mill at a frequency of 30 Hz for 30 seconds. DNA samples were eluted in 50-100 µl DES water and DNA concentrations were determined using the Qubit™ dsDNA HS Assay Kit (Invitrogen, life technologies) on Qubit®2.0 (Life Technologies).

### Amplicon generation and library construction

16S rRNA amplicon generation and library preparation were performed according to Liu et *al*.(33). Amplicon libraries were generated using the PCR primers 799F (AACMGGATTAGATACCCKG)(24), and 1193R (5’-ACGTCATCCCCACCTTCC-3’)(23). The first amplification was performed in a 25 µL reaction volume, including 2.5 µl microbial DNA (5ng/µl), 5 µl of 799F-B primer (1µM), 5µl of 1193R-B primer (1µM), 12.5µl 2X KAPA HiFi Hot Start Ready Mix. The PCR setting was 95°C for 3 minutes, 30 cycles of 95°C for 30 seconds, 55°C for 30 seconds, 72°C for 30 seconds, and 72°C for 5 minutes. The second amplification was conducted in 20 µL reaction volume, each containing 10 µL 2x KAPA HiFi Hot Start Ready Mix, 200 nM of 2P-F and 2P-R primer, 2 nM of F-(N) and R-(N) primer, and 40ng first-round PCR product. The PCR conditions were 95°C for 3 minutes, 10 cycles of 95°C for 30 seconds, 55°C for 30 seconds, 72°C for 30 seconds, and 72°C for 2 minutes. All the primers sequences were included in the Table S1.

After each PCR, the PCR products were loaded on 2% agarose gel and cut from the gel with Tanon UV-2000 gel imaging system and extracted from the agarose using the QIAquick Gel Extraction kit (Qiagen). DNA concentrations were determined using the Qubit™ dsDNA HS Assay Kit on Qubit®2.0.

After two-round PCR, 5 biological replicates for each genotype were combined, and DNA concentrations were determined using the Qubit™ dsDNA HS Assay Kit on Qubit®2.0. Then all the genotype samples were combined with same amount of each genotype PCR product.

### Sequencing and data analysis

The libraries were sent to Novogene company for 16s rRNA gene sequencing. Processing and statistical analysis of 16S rRNA counts was performed by the Core Facility of Bioinformatics in Shanghai Center for Plant Stress Biology, China.

The quality of reads was checked with fastqc v 0.11.7 and the data was pre-processed with trimmomatics v0.39 (34). The processed high-quality data was assembled with FLASH v1.2.11(35). QIIME (version 1.9.1) was mainly used for subsequent analysis(36). Assembled reads were demultiplexed with *split_library_fastq*.*py* and chimeric sequences were removed by *identify_chimera_seqs*.*py* with usearch (-m usearch61) method and “Gold” database (-r gold.fa). Open-reference OTU picking was carried out using the *pick_open_reference_otus*.*py* script in QIIME at 97% sequence identity with GreenGenes 16S database v.13_8. After removing OTUs which belong to mitochondria, Chlorophyta, Archaea and Cyanobacteria, we got 6275 OTUs. The beta-diversity analysis and statistical analysis were performed mainly according to Schlaeppi et. *al* (28). We generated the rarefied tables (100x tables from 700-70000 sequences per sample, step of 700 sequences), selected #50 of rarefied OTU tables as the threshold-independent community (TIC) table. A minimum of 20 sequences per OTU in at least one sample was used as a criterion to define Abundant Community members (ACM)(28). The relative abundance of ACM was calculated by dividing the reads per ACM in a sample by the sum of the usable reads in that sample. Pairwise UniFrac distance and principal coordinates analysis were performed by QIIME. The significant differences between samples were assessed by the ANOVA-based statistics. The resulting p-values were adjusted by Benjamini-Hochberg false discovery rate (FDR) correction for multiple hypotheses testing.

---

**Figure S1.**
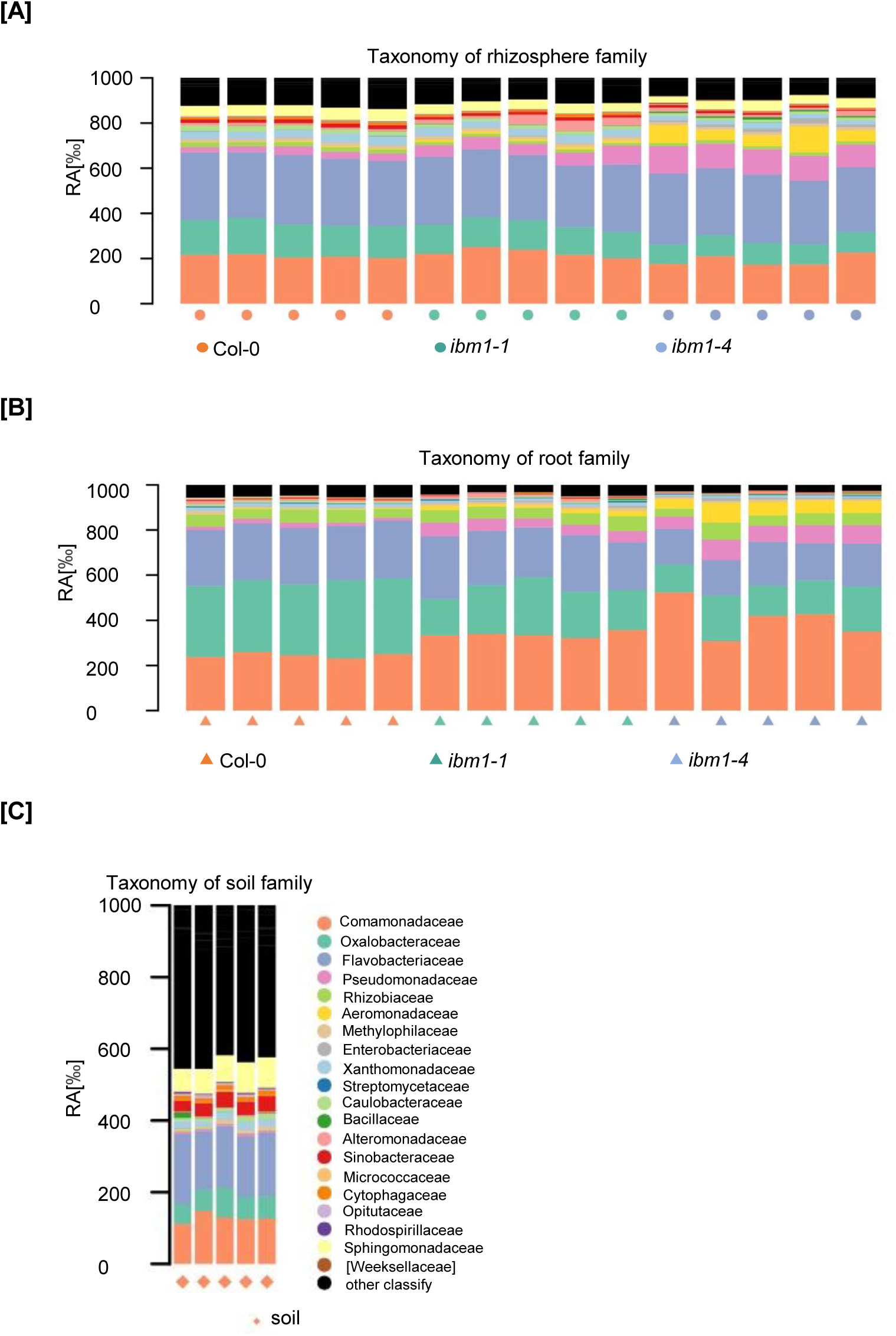
Taxonomic structure at the family rank of the ACM in Col-0, *ibm1-1* and *ibm1-4* at rhizosphere **[A]**, endosphere **[B]** and soil sample **[C]**.

**Figure S2.**
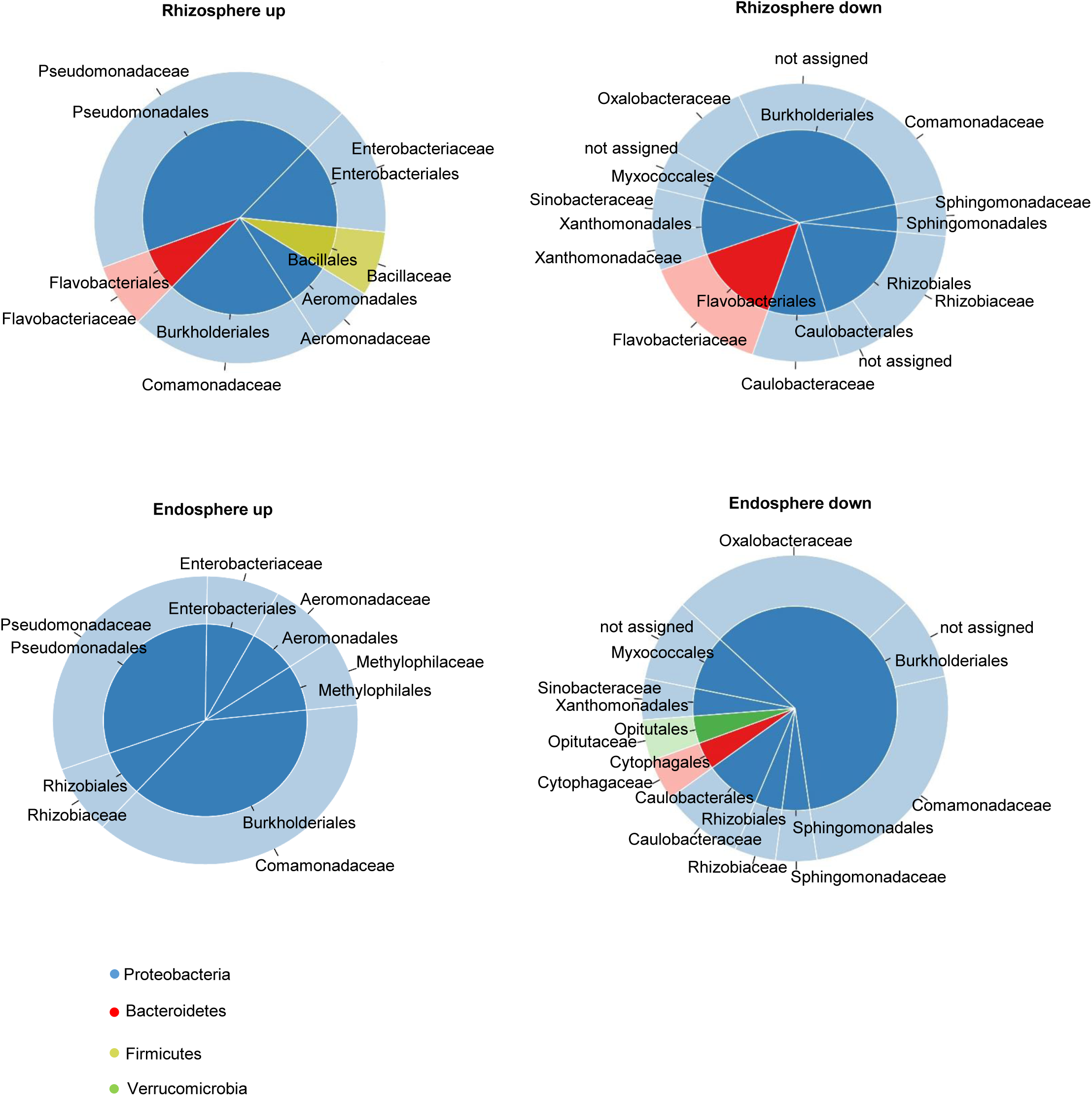
The composition of the ACMs which are commonly affected by *ibm1-1* and *ibm1-4* (p<0.05). The color in the pie chart represents the bacterial phyla of the corresponding taxa. The ACMs were displayed at order and family rank in the inner and the outer ring of the pie chart, respectively.

**Table S1. Primers used in this study**

